# Potential Pharmacodynamic Mechanism of the Main ingredients in Licorice for Chronic Obstructive Pulmonary Disease

**DOI:** 10.1101/2021.08.29.458060

**Authors:** Cai Chen, Jianpeng An, Guodong Shen, Yang Shen

**Affiliations:** Shandong Institute of Advanced Technology Chinese Academy of Sciences, Jinan, 250000, China; Integrated Traditional Chinese and Western Medicine Hospitial of Tongzhou District, Beijing, China; Ministry of Environmental Protection, School of Public Health, Tongji Medical College, Huazhong University of Science and Technology, #13 Hangkong Road, Wuhan, 430030, Hubei, China

**Author notes:** These authors contributed equally to this work (Cai Chen, Jianpeng An). Corresponding author: YangShen, Ministry of Environmental Protection, School of Public Health, Tongji Medical College, Huazhong University of Science and Technology, #13 Hangkong Road, Wuhan, 430030, Hubei, China.

**Keywords:** COPD, licorice, Gene Ontology, Kyoto Encyclopedia of Genes and Genomes, network pharmacology

## Abstract

**Purpose:** This study aimed to investigate the effect of essential ingredients of licorice on the chronic obstructive pulmonary disease (COPD).

**Method:** The ingredients information were obtained from *PubChem* (https://pubchem.ncbi.nlm.nih.gov/), related genes about COPD was collected from *geneCards* (http://www.genecards.org/). *Network* pharmacology was utilized in this study.

**Result:** The intersection data set contains 20 molecular targets between COPD and liquorice. Protein-protein interaction network showed that there are a total of 58 nodes and 137 edges involved. The link number of AKT1 in PPI network was 39, which is the highest level of interaction. MAPK1 is an important target of Licorice on COPD.

**Conclusion:** MAPK signaling pathway could be the important key target of main ingredients of licorice on COPD.

## I. Introduction

Chronic obstructive pulmonary disease (COPD) is one of the common preventable and treatable disease, with the emblematic symptom of persistent airflow restriction, which is associated with an increased chronic inflammatory response of the airways and lungs to toxic particles or gases[1-4]. According to the World Health Organization, approximately 3 million people in the world dead as a consequence of COPD each year. One meta-analysis of global prevalence of COPD, which purposed to examine the global prevalence of COPD in men and women, revealed that prevalence rate of COPD was 9.23% (95% CI:8.16%-10.36%) among male group and 6.16% (95% CI:5.41%-6.95%) among female group[5]. Moreover, there were some complications of COPD, such as cardiovascular disease, anxiety and depression, lung cancer [6-8], which leads to the lower living quality and much more burden of patient ‘s life[9, 10].

Liquorice, widely distributed among Asia, Europe, America, etc., has been used as medicinal material for long years[11-13]. Some researches have proved that liquorice has the function of anti-inflammation, preventing coughing, antianaphylaxis and so on[14, 15]. It was reported that the essential ingredients of liquorice are glycyrrhizic acid, glycyrrhizin, quercetin, and formononetin[16, 17]. Generally, liquorice, the whole plant is utilized to made as pills or decoction. Consequently, the mechanism and molecular pathways for every component was not clear. Up to now, much researches concentrated on the biological effect of liquorice as a whole, while the molecular effect of its every component hasn’t been figured out. Considering its bargain price, widespread distribution and unique biological effect, this research purposed to investigate the mechanism of essential component’s signaling pathways (formononetin, liquiritin, glycyrrhizic acid, glabridin, quercetin and isoliquiritigenin) based on network pharmacology.

## II. Method and Materials

The ingredients of liquorice involved in this research were formononetin, liquiritin, glycyrrhizic acid, glabridin, quercetin and isoliquiritigenin, respectively. Their two-demension and three-demension struction were from public database, *PubChem* (https://pubchem.ncbi.nlm.nih.gov/). Pharmacokinetic information of these compound was obtained from *Traditional Chinese Medicine Systems Pharmacology (TCMSP)*, including drug-likeness (DL), oral bioavailability (OB), intestinal epithelial permeability (Caco-2) and number of H-bond donor/acceptor (Hdon/Hacc).

The target molecule of the Liquorice ingredients were attained from public database *Swisstargetprediction* (http://www.swisstargetprediction.ch/). The potential molecular target was selected when the probability was above zero. The related genes about COPD was collected from *geneCards* (http://www.genecards.org/), and the genes were used in this research when *relevance score* was more than 30.

Construction and analysis of component-action target network was based on Cytoscape software (version 3.7.2). Six compound and common target between COPD and Liquorice was imported to Cytoscape, after position adjustment network was finished.

The String database (https://string-db.org/) is one database containing known and predicted protein-protein interactions, in which a large number of protein-protein interactions were collected, involving a total of 9643763 proteins and 138838440 interactions, including data detected experimentally and predicted by bio-information methods. The common targets between disease and Liquorice were imported into the String database to define human species and obtain the protein interaction relationship. The results were saved in TSV format. The node1, node2 and combined score information in the file was retained and imported into Cytoscape software to draw the interaction network. The core of protein-protein interaction network was calculated in R software. Then GO enrichment analysis and KEGG pathway annotation analysis were finished in *Rstudio* with the p-value <0.05 as well.

## III. Result

***Figure 1*** and ***Figure 2*** showed the two and three dimension structure of formononetin, liquiritin, glycyrrhizin, glabridin, quercetin and isoliquiritigenin, respectively. Pharmacokinetic information was illustrated in ***Table 1***. The number of H-bond donor/acceptor for formononetin, liquiritin, glycyrrhizin, glabridin, quercetin and isoliquiritigenin were 1/4, 8/16, 5/9, 2/4, 5/7 and 6/9, respectively. The oral bioavailability of formononetin, liquiritin, glycyrrhizin, glabridin, quercetin and isoliquiritigenin were 69.67, 9.06, 65.69, 53.25, 46.43 and 8.61, respectively.

**Figure 1.**
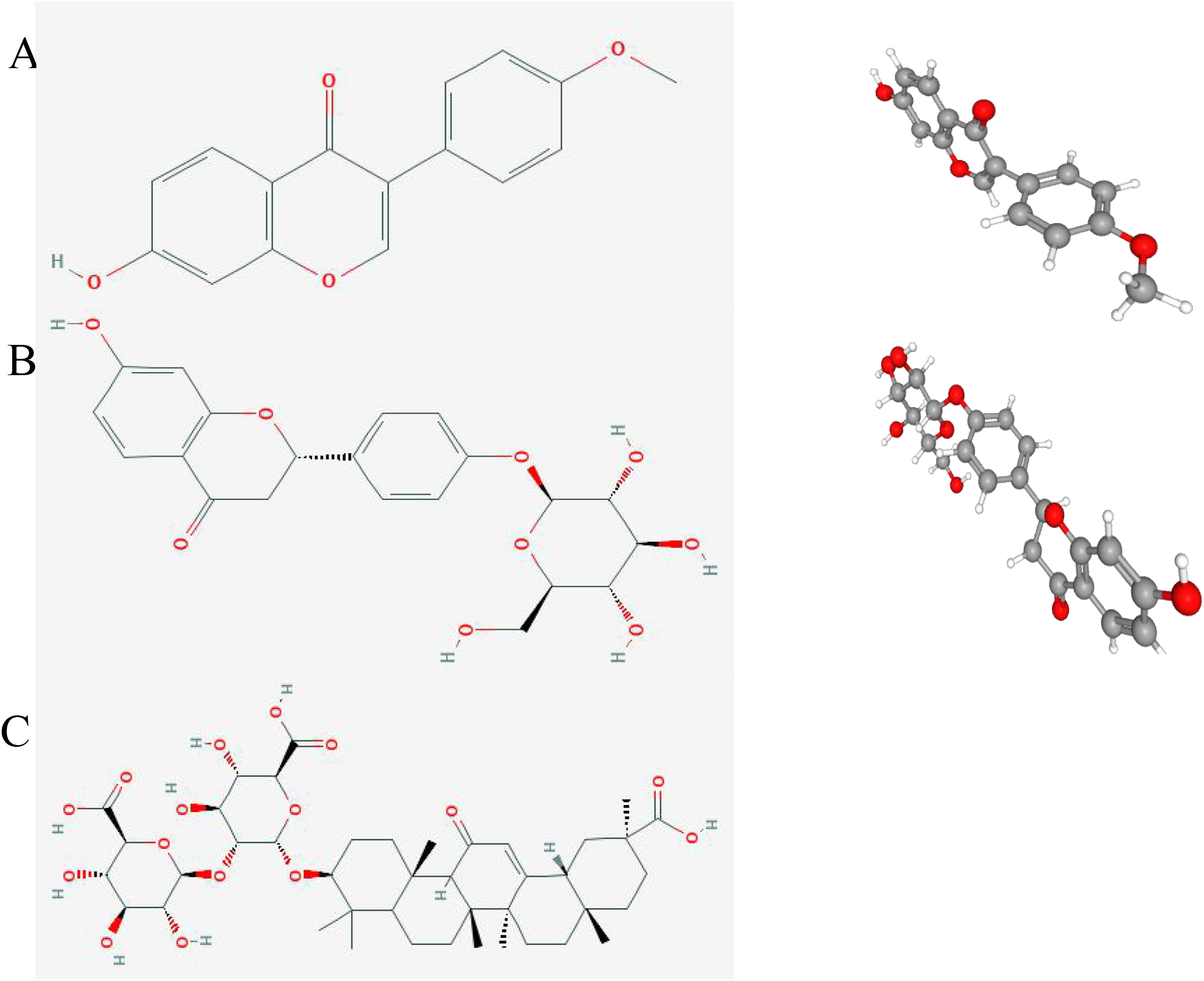
The two and three demonsion structure compounds (A: formononetin; B: liquiritin; C: glycyrrhizin)

**Figure 2.**
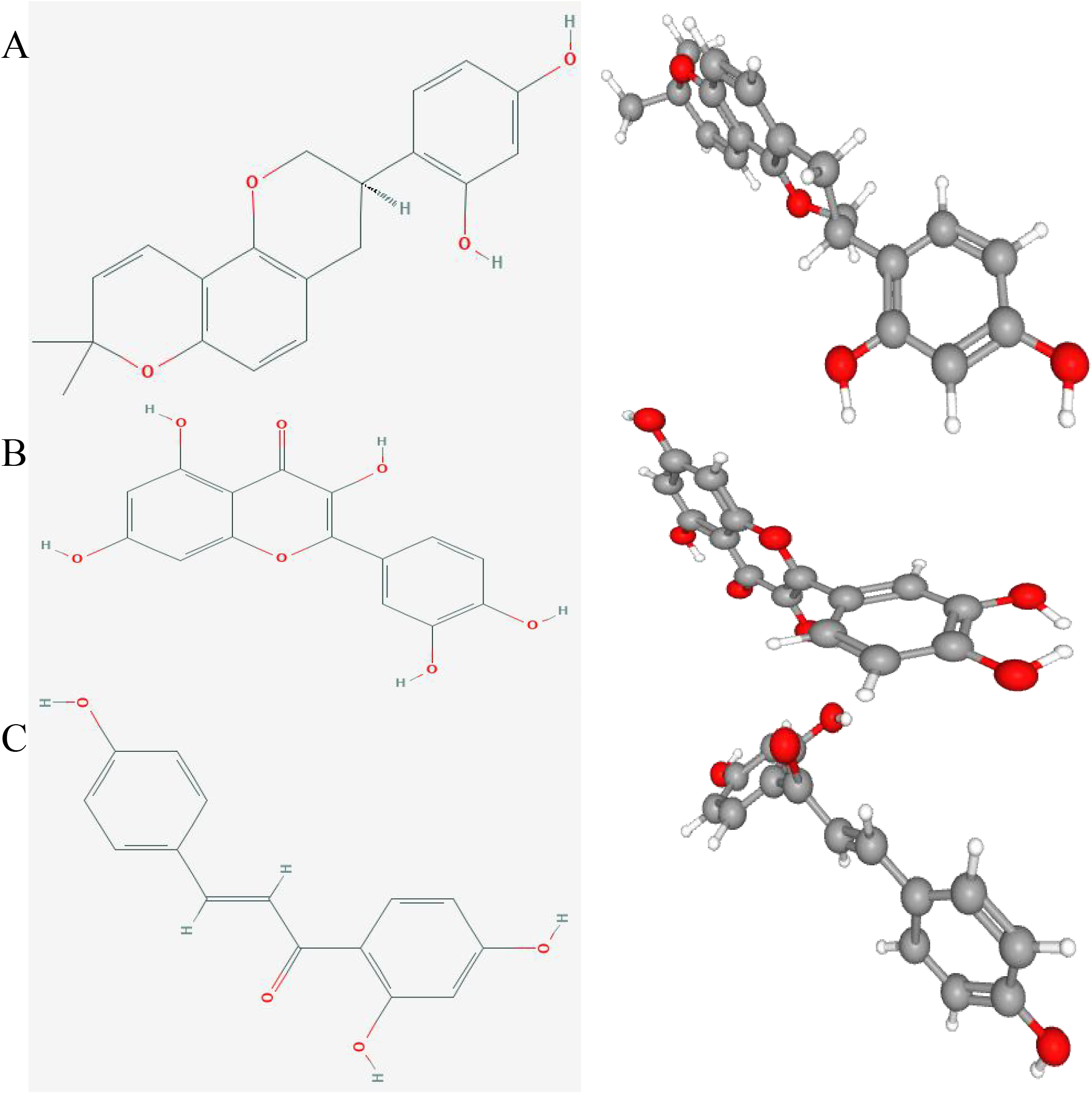
The two and three demonsion structure of conpounds (A: glabridin; B: quercetin; C: isoliquiritigenin)

**Table 1.** Pharmacokinetic information about six compounds

|  | Hdon | Hacc | OB (%) | Caco-2 | DL |
| --- | --- | --- | --- | --- | --- |
| Formononetin | 1 | 4 | 69.67 | 0.78 | 0.21 |
| Liquiritin | 8 | 16 | 9.06 | -2.23 | 0.11 |
| Glycyrrhizin | 5 | 9 | 65.69 | -1.06 | 0.74 |
| Glabridin | 2 | 4 | 53.25 | 0.97 | 0.47 |
| Quercetin | 5 | 7 | 46.43 | 0.05 | 0.28 |
| Isoliquiritigenin | 6 | 9 | 8.61 | -1.36 | 0.6 |

There were 417 target molecule for COPD according to relevance score mentioned above and 64 target molecule for Liquorice ***(Figure 3)***. The intersection data set contains 20 molecular targets between COPD and liquorice. The information of the six active ingredientsand molecular targets of Licorice was introduced into Cytoscape to construct the network, as shown in ***Figure 4***. There are a total of 58 nodes and 137 edges involved. The black type indicates the six ingredients of Licorice, blue ellipse represented the potential target. It can be seen from the ***Figure 4*** that the same target could be corresponding to different active ingredients or the same active ingredient, which fully reflects the multi-component and multi-target action characteristics of Liquorice.

**Figure 3.**
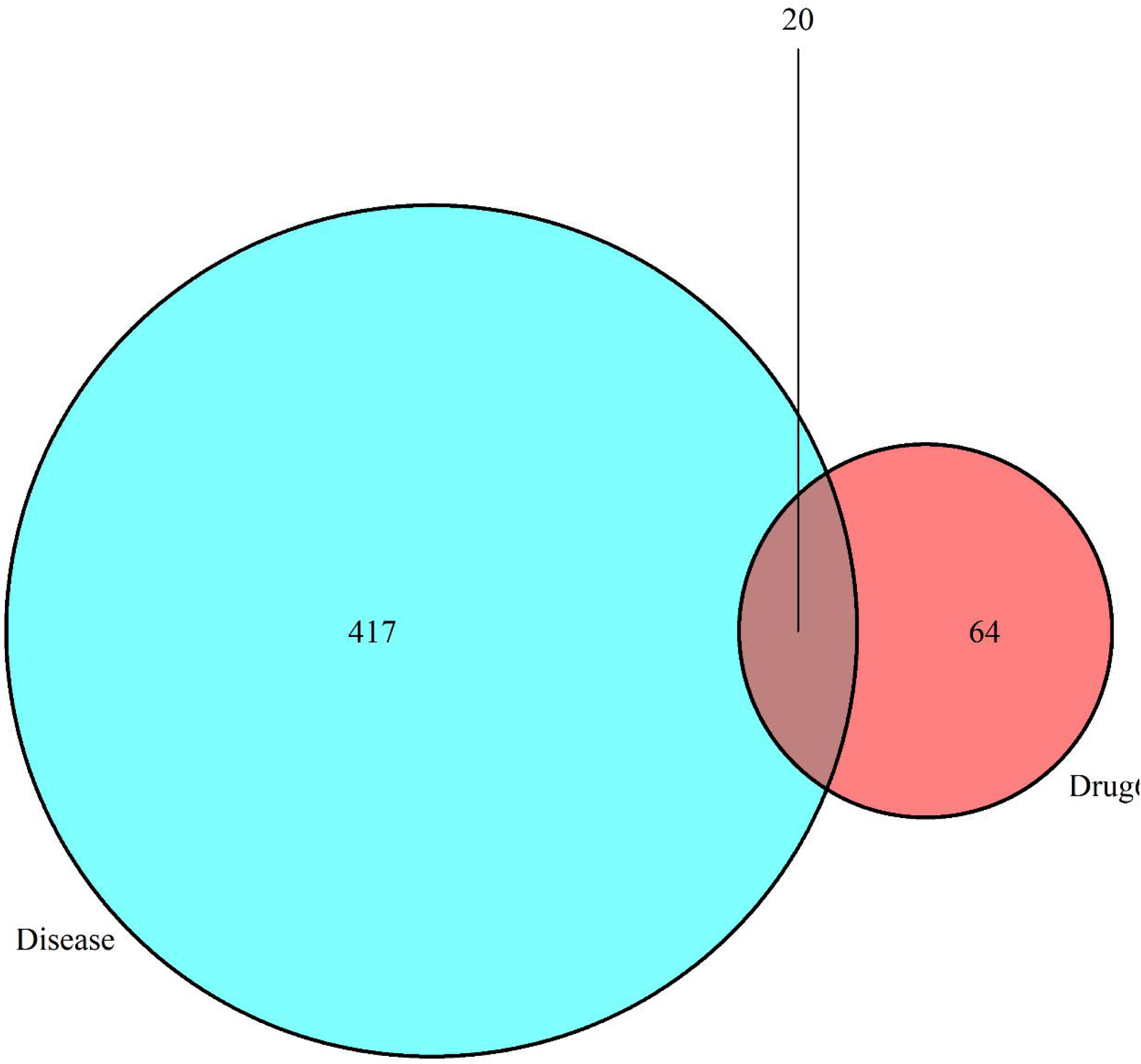
The target molecule of COPD and liquorice

**Figure 4.**
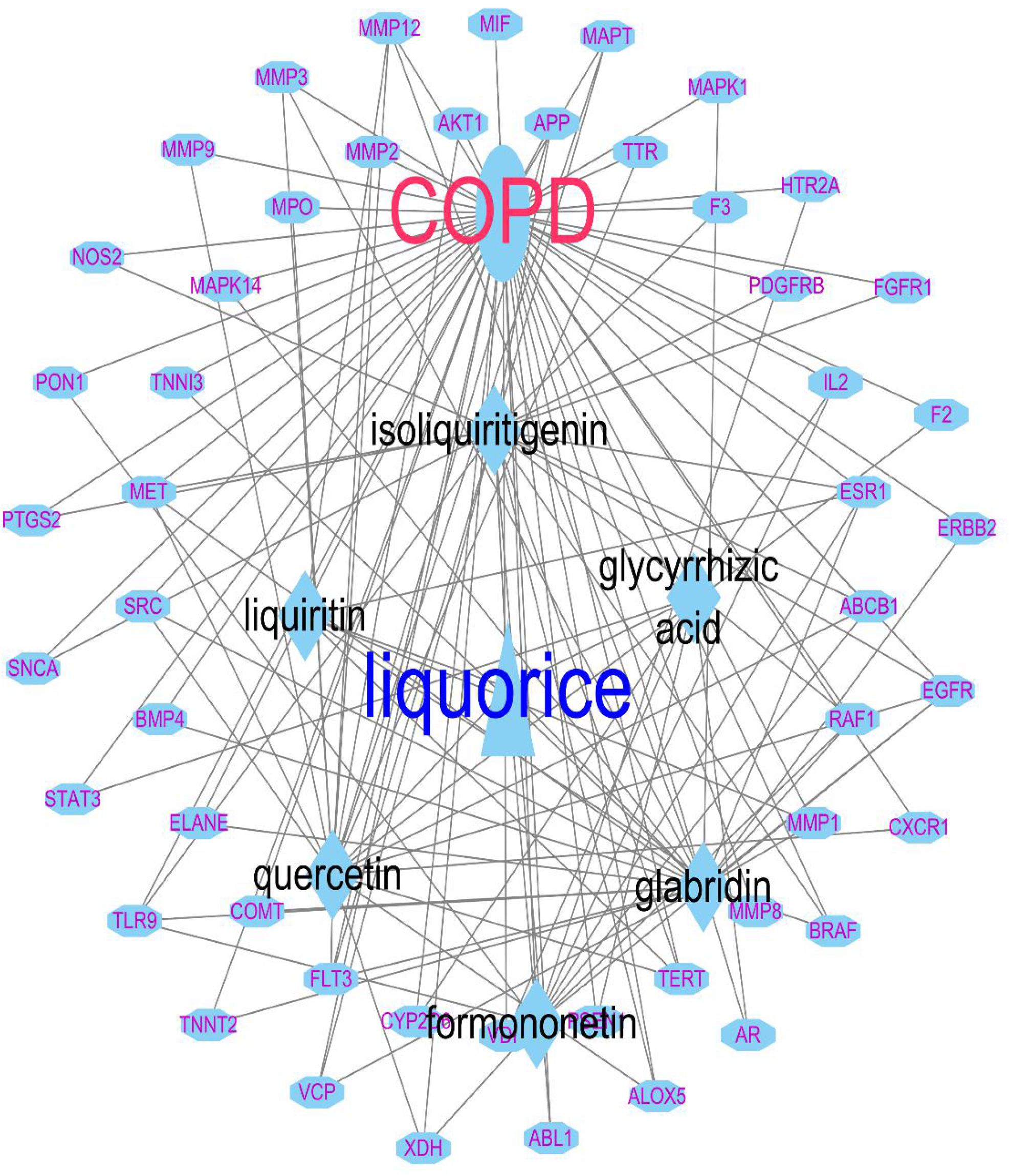
The interaction network COPD, liquorice and molecular targets

***Figure 5*** was the result after the process of String database. Color ‘green’ is the gene neighborhood, ‘black’ represents the co-expression, ‘blue’ is gene co-occurrence, ‘red’ is the gene fusions. As shown in ***Figure 6***, the link number of AKT1 was 39, which is the highest level of interaction. ***Figure 7*** presented the top ten key targets. GO enrichment analysis is a finite acyclic graph that counts the number or composition of proteins or genes at a functional level. The generatio of endopeptidase activity and phosphatase were greater than the others. Nuclear receptor activity and transcription factor activity, direct legend regulated sequence specific DNA binding were least (***Figure 8,Table 2***). The results of KEGG analysis are shown in ***Figure 9***. The counts of proteoglycans in cancer, endocrine resistance, MAPK signaling pathway, EGFR tyrosine kinase inhibitor resistance, relaxin signaling pathway, Rap1 signaling pathway was 14,12, 12,10,10 and 10, respectively.

**Figure 5.**
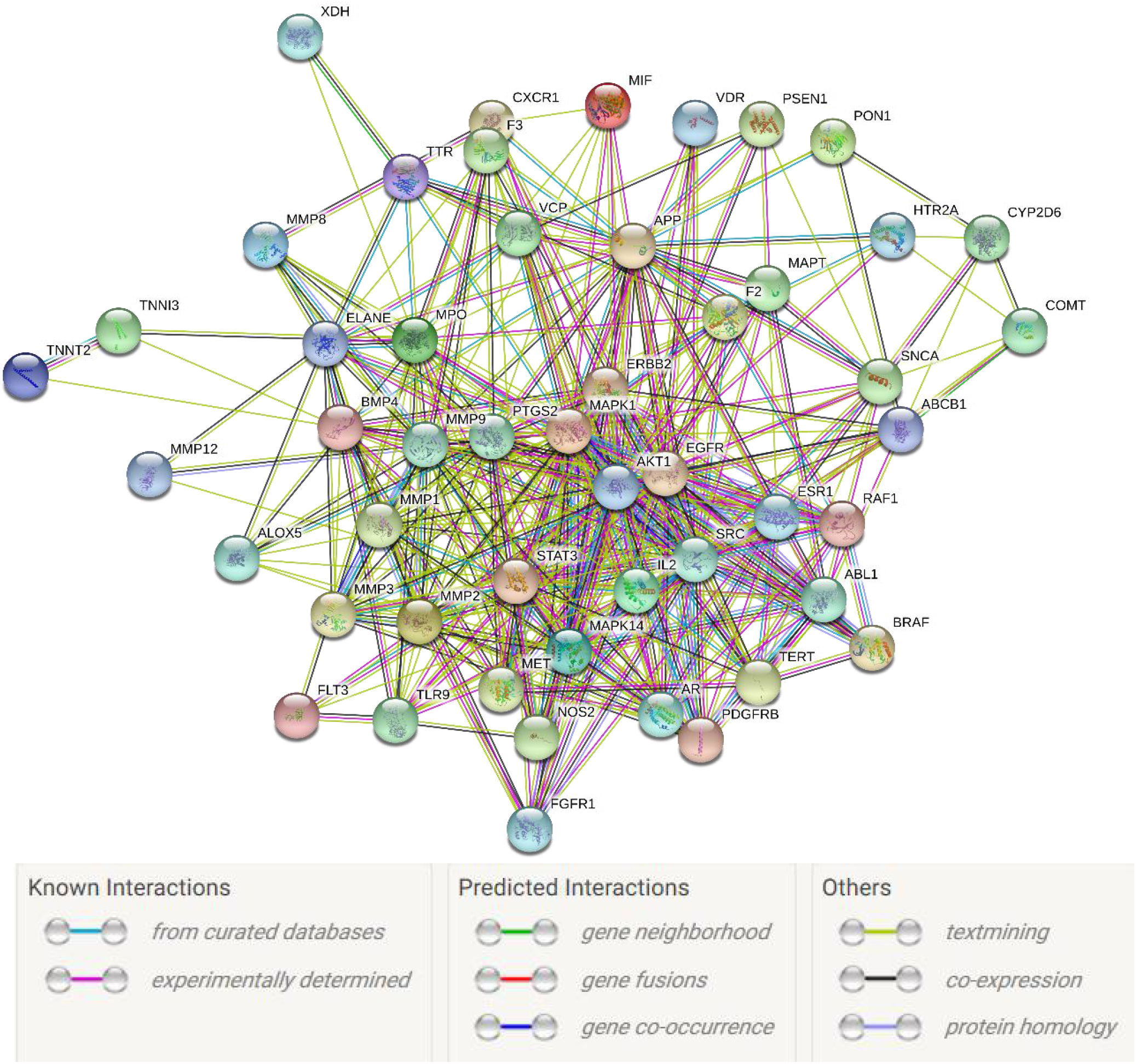
The structure of protain-protain interaction network

**Figure 6.**
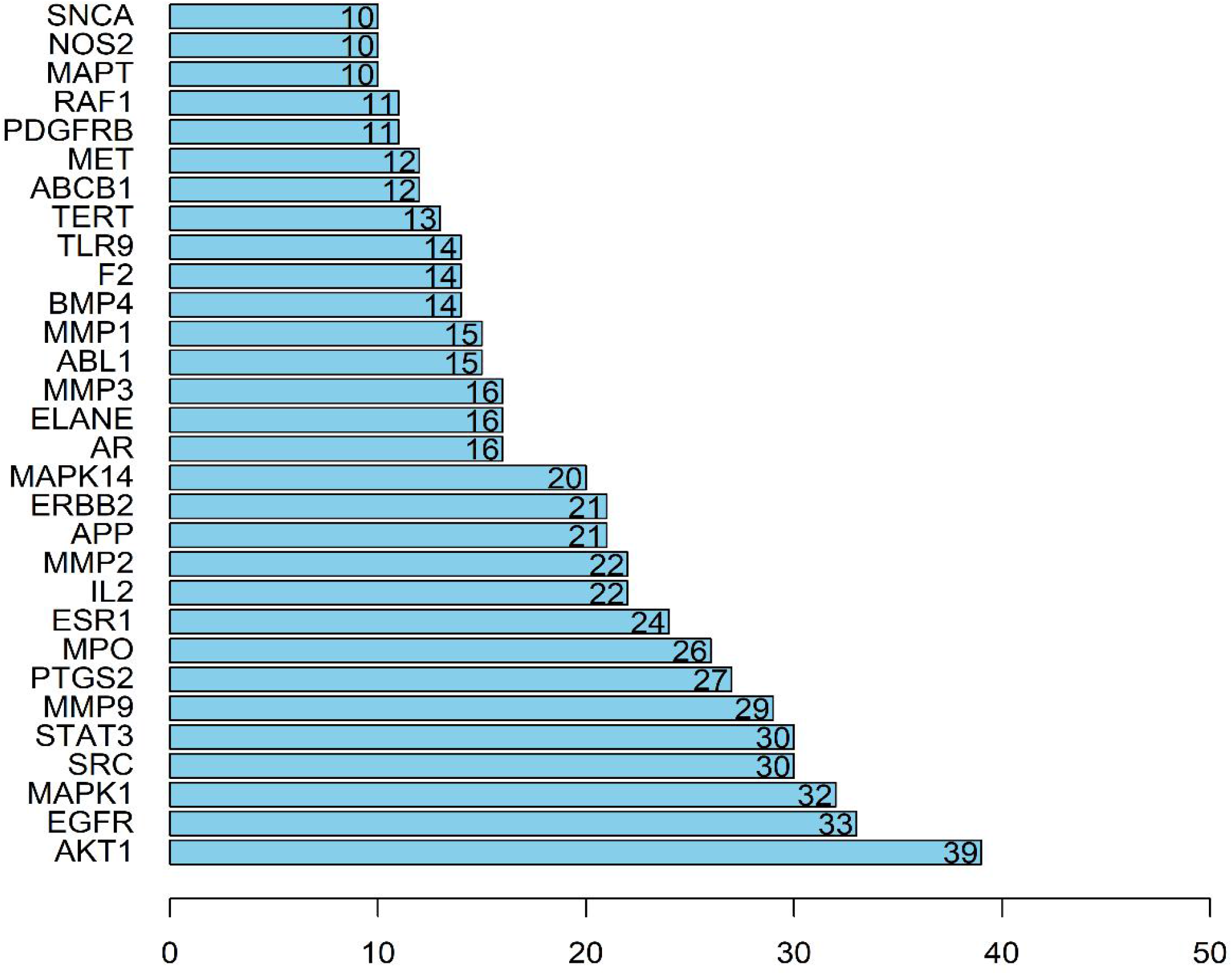
PPI network link number

**Figure 7.**
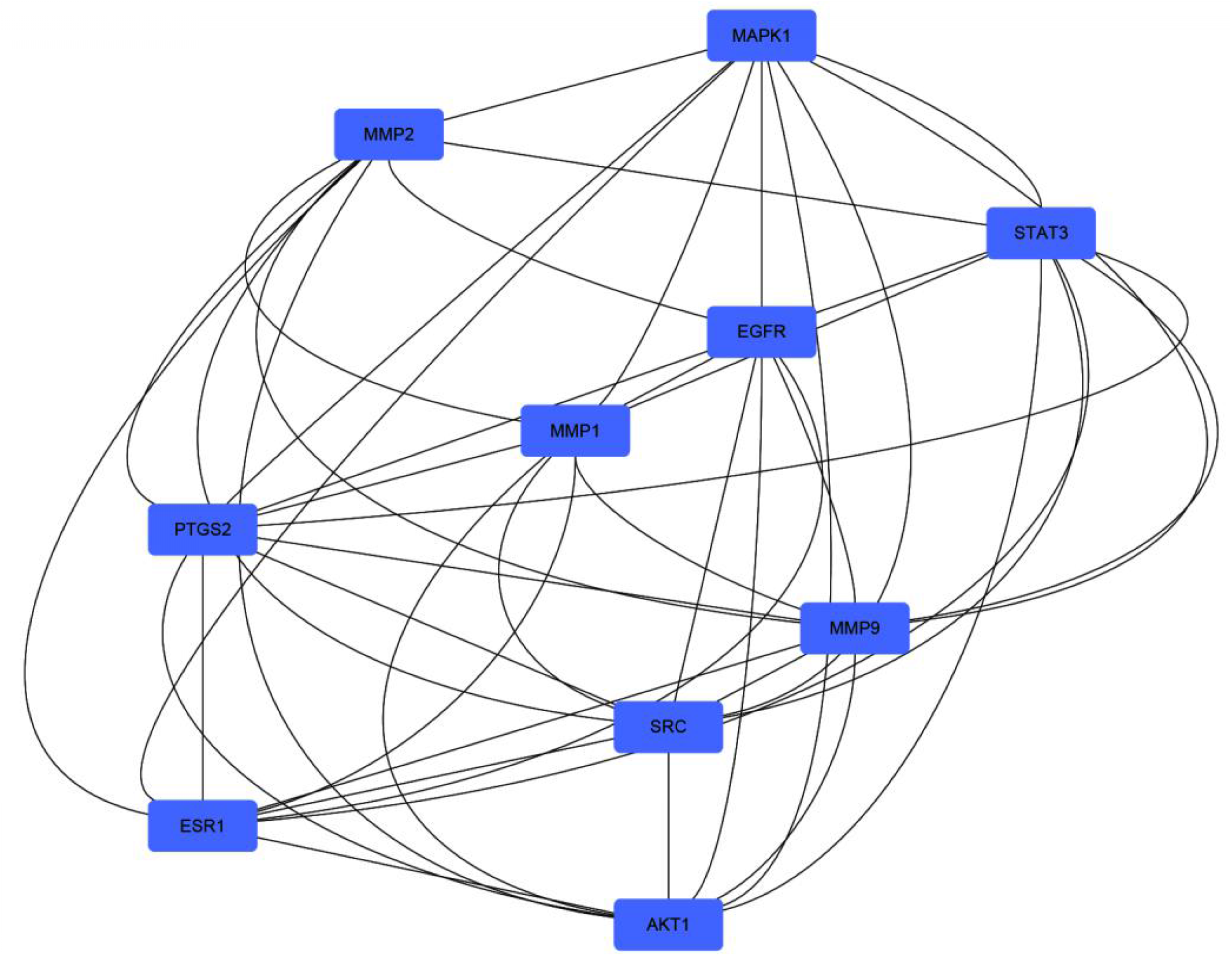
The top ten targets in the PPI network

**Figure 8.**
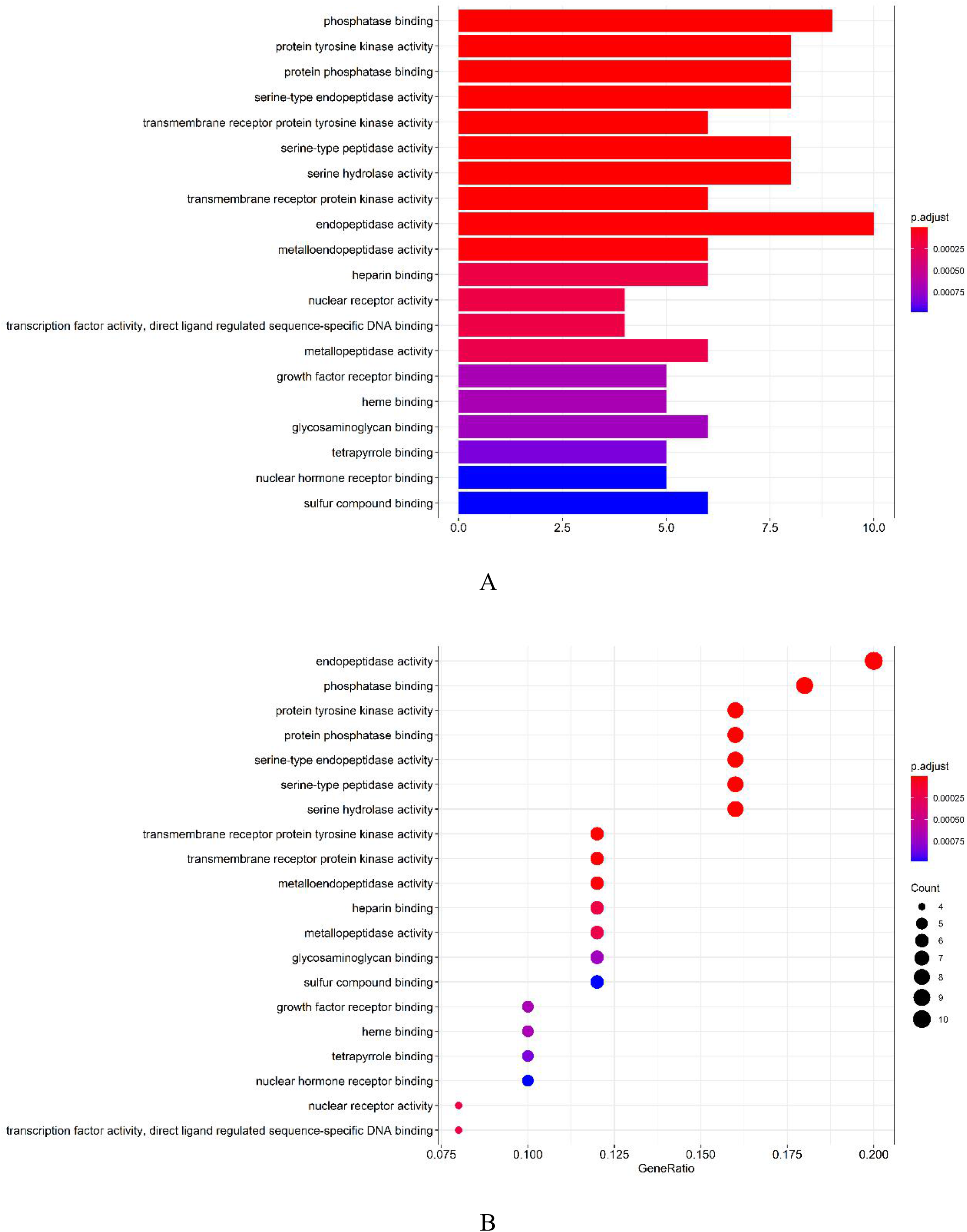
The result of GO enrichment analysis

**Table 2.**
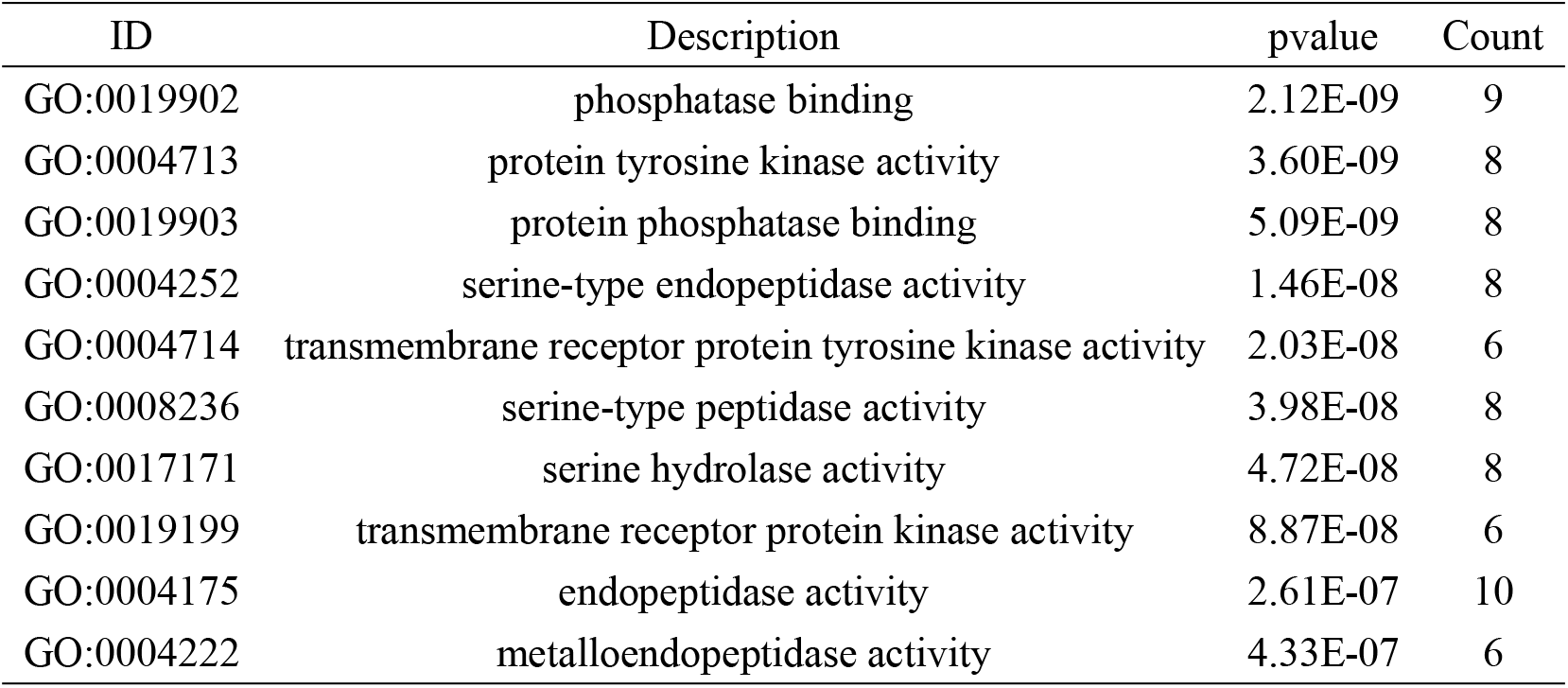
GO enrichment analysis result

**Figure 9.**
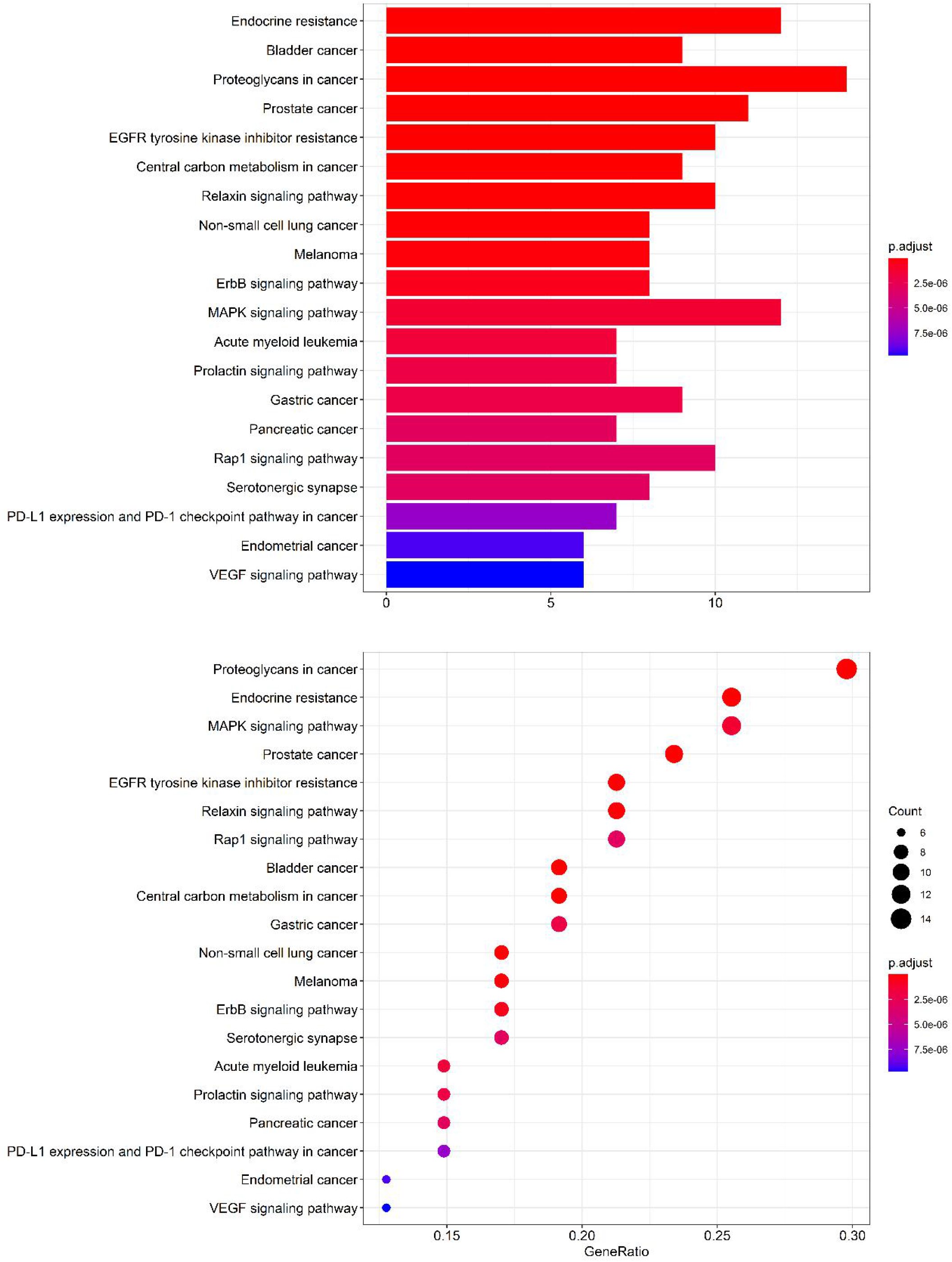
The result of KEGG enrichment analysis

## IV. Discussion

COPD is a chronic bronchitis and/or emphysema characterized by airflow obstruction that could progress to pulmonary-heart disease and respiratory failure as a common chronic disease[2, 4, 18]. It has been formed a broad consensus that COPD is associated with abnormal inflammatory reactions with high morbidity and mortality[19]. Licorice has been internationally utilized for medicinal herb, while its mechanism of some main ingredients for COPD still kept unclear. The purpose of the present study was to investigate the effect of main ingredients on COPD according to network pharmacology.

In this study, it revealed that Akt1 is the common target between COPD and licorice. AKT1, also named as protein kinase B, is one of 3 closely related serine/threonine-protein kinases (AKT1, AKT2 and AKT3) called the AKT kinase, and which regulates many processes including metabolism, proliferation, cell survival, growth and angiogenesis[20-22]. It appeared as one key node in PI3K-AKT signaling, protects against acute lung injury[23]. Qu ‘s experiment indicated that glycyrrhizic acid inhibited the production of inflammatory factors in LPS-induced ALI by regulating the PI3K/AKT/mTOR pathway related autophagy[24]. Vito Lorusso and Ilaria Marech summarized that isoliquiritigenin could suppress HIF-1α level, VEGF expression and secretion, cell migration and to reduce the expression and secretion of MMP-9/-2 and these effects might be mediated through inhibition of p38, PI3K/Akt and NF-κB signaling pathways[25].

As shown in ***Figure 8***, MAPK1 was one of critical targets, which is an important transmitter of signals from the cell surface to the inside of the nucleus. The result of KEGG enrichment analysis also illustrated that licorice relieves COPD symptoms via MAPK signaling pathway. It has been reported that MAPK pathway is one of the common intersection pathways of signal transduction pathways including stress, inflammation, cell proliferation, differentiation, functional synchronization, transformation, apoptosis and so on. Previous studies demonstrated that MAPK signal pathway was involved in the inflammation reaction and oxidative stress[26, 27]. Some researchers also found that the ingredients of licorice has the function to mediate the expression of MAPK signaling pathway[28, 29].

The limitation of the present study was that this study was performed abstractly, in the future study, we will operate animal experiment to prove it with ethical approval.

## V. Conclusion

MAPK signaling pathway could be the important key target of main ingredients of licorice on COPD.

## VI. Availability of data and materials

The ingredients of liquorice involved were attained from *PubChem* (https://pubchem.ncbi.nlm.nih.gov/). Pharmacokinetic information of these compound was obtained from *Traditional Chinese Medicine Systems Pharmacology (TCMSP)*.

The target molecule of the Liquorice ingredients were attained from public database *Swisstargetprediction* (http://www.swisstargetprediction.ch/). The related genes about COPD was collected from *geneCards* (http://www.genecards.org/).

## VII. onsent to Publish

All author consent to publish this article in this article.

## Acknowledgement

This study was supported by Project of Jinan “20 Universities” (#2019GXRC040).

